# A murine model of *Trypanosoma brucei-*induced myocarditis and cardiac dysfunction

**DOI:** 10.1101/2023.10.05.560950

**Authors:** Nathan P. Crilly, Marcelle Dina Zita, Alexander K. Beaver, Polina Sysa-Shah, Aashik Bhalodia, Kathy Gabrielson, Luigi Adamo, Monica R. Mugnier

**Author notes:** Corresponding authors: Monica R. Mugnier.; Luigi Adamo. Co-Senior authors (Luigi Adamo, Monica R. Mugnier). **Conflict-of-interest statement:** The authors have declared that no conflicts of interest exist.

## Abstract

*Trypanosoma brucei* is a protozoan parasite that causes human and animal African trypanosomiases (HAT and AAT). Cardiac symptoms are commonly reported in HAT patients, and intracardiac parasites with accompanying myocarditis have been observed in both natural hosts and animal models of *T. brucei* infection. Despite the importance of *T. brucei* as a cause of cardiac dysfunction and the dramatic socioeconomic impact of African trypanosomiases in sub-Saharan Africa, there are currently no reproducible murine models of *T. brucei-*associated cardiomyopathy. We present the first clinically relevant, reproducible murine model of cardiac dysfunction in chronic *T. brucei* infection. Similar to humans, mice showed histological evidence of myocarditis and elevation of serum NT-proBNP with electrocardiographic abnormalities. Serum NT-proBNP levels were elevated prior to the development of severe ventricular dysfunction. On flow cytometry, myocarditis was associated with an increase of most myocardial immune cell populations, including multiple T cell and macrophage subsets, corroborating the notion that *T. brucei-*associated cardiac damage is an immune-mediated event. This novel mouse model represents a powerful and practical tool to investigate the pathogenesis of *T. brucei*-mediated heart damage and supports the development of therapeutic options for *T. brucei*-associated cardiac disease.

## Introduction

The threat of *Trypanosoma brucei* is a fact of life throughout sub-Saharan Africa. A protozoan parasite spread by the tsetse fly vector, *T. brucei* is the causative agent of two neglected tropical diseases, human and animal African trypanosomiasis (HAT and AAT) (1). In both humans and animals, the disease is characterized by waxing and waning fever followed by chronic wasting and a coma which gives the disease its historical name, “sleeping sickness” (2, 3). Two subspecies are responsible for human infection: *T. b. gambiense*, which is endemic in West and Central Africa and *T. b. rhodesiense,* which is restricted to East and Southern Africa (4). The disease character and progression are highly variable, although *T. b. rhodesiense* infection is generally aggressive and rapidly progressive, while *T. b. gambiense* infection tends to follow a more chronic, slowly progressive course (2). Although HAT is uncommon, with fewer than 1000 cases reported in 2019 through 2020, the ubiquity of animal reservoirs and the insect vector throughout sub-Saharan Africa means that re-emergence is a constant threat (5–8). In addition to the direct impacts on human health, AAT is a severe burden to animal agriculture throughout sub-Saharan Africa and a serious contributor to food insecurity in the region, with an estimated cost of $4.75 billion per year and an incalculable impact on individual livelihoods (9).

*T. brucei* is often thought of as a parasite of the hemolymphatics and central nervous system. However, both historical and recent studies have led to a renewed appreciation for *T. brucei* as a parasite of extravascular spaces (10, 11). *T. brucei* parasites invade and colonize multiple extravascular tissues, including the heart, lungs, adipose tissue, skin, eyes, gonads, pancreas, and brain (12, 13). Within these spaces, tissue-resident parasites are associated with widespread inflammation and damage, resulting in loss of organ function that contributes to mortality (10, 11, 14).

*T. brucei* colonizes the heart, and cardiac disease is a likely contributor to HAT mortality (15, 16). HAT patients exhibit elevated cardiac NT-proBNP, a biomarker which is a sensitive predictor of heart failure, and 19% report at least one symptom consistent with heart failure (16). Although electrocardiography (ECG) is not routinely performed on HAT patients, electrocardiographic abnormalities have been reported in up to 71% of HAT patients, with most notable changes including prolonged QT interval and low voltage(16, 17). One study found that patients in Cameroon with dilated cardiomyopathy were significantly more likely to have anti-trypanosomal antibodies than healthy controls, suggesting that even clinically inapparent *T. brucei* infection might contribute to cardiac dysfunction(18). In agreement with these clinical findings, myocarditis has been reported in autopsies of patients who have died from HAT, suggesting that cardiac inflammation is a contributor to *T. brucei*-associated cardiac disease (19, 20). Cardiac symptoms and resulting pathology have been reported in both *T. b. gambiense* and *T. b. rhodesiense* infection (16). There is some evidence that *T. b. rhodesiense* causes more severe cardiac pathology, but the small number of recorded cases make evaluation difficult (19, 21).

Unfortunately, while there is clear evidence that cardiac disease is a consequence of African trypanosomiasis., *T. brucei*-associated heart disease remains understudied. In particular, there has been a lack of well-established animal models to allow detailed investigation into the pathology, pathogenesis, and functional consequences of *T. brucei*-associated heart disease. Several historical studies have described the histopathology of myocarditis during *T. brucei* infection, especially in large animal models such as cattle, dogs, and nonhuman primates (22–25). While large animal models, especially primates, accurately recapitulate human disease, the expense, difficulty, and ethical issues of working with such model species has prevented their widespread adoption by *T. brucei* researchers. More recently, researchers have published a rat model of acute cardiac disease during early *T. brucei* infection. In this acute model, infected rats exhibited myocarditis and ventricular arrhythmias at 11 days postinfection (dpi), similar to the pathological and EKG findings reported in human patients (16, 20, 26, 27). Although the rat model demonstrates that *T. brucei*-related cardiac disease can be recapitulated in a rodent, there are important limitations. Notably, studies in the rat have been limited to the first two weeks of infection, while clinical data suggest that cardiac symptoms and myocarditis are most prominent during chronic HAT (19, 20). In addition, studies in the rat model have been limited to analyzing histopathology and ECG findings, although other modalities such as echocardiography and measurement of cardiac biomarkers are important for assessing the severity and progression of heart disease. Finally, there are relatively few tools for genetic and immunological manipulations in the rat, potentially limiting its usefulness for investigating disease pathogenesis (28) .

A mouse model of *T. brucei*-associated cardiac disease would have multiple advantages. Compared to other animal models, mice are relatively inexpensive, safe, and simple to handle and house, making them practical for most investigators. In addition, there are numerous well-characterized strains and knockout lines of mice, allowing detailed investigation into disease mechanisms (28). Mice are also well-characterized as a model species for a variety of parasitic diseases, including those caused by African (*T. brucei*) and New World (*Trypanosoma cruzi*) trypanosomes (27, 28). Historical studies performed with infection protocols that cannot be replicated in modern times showed evidence of myocarditis in *T. brucei*-infected mice via histopathology, further supporting the utility of the mouse as a model of *T. brucei*-associated cardiac disease, although the consequences of this inflammation were not evaluated (29, 30). Overall, the establishment of a robust, reproducible mouse model would allow for a detailed investigation into the mechanisms and consequences of *T. brucei*-associated cardiac disease and would be instrumental to the development of novel therapeutic options.

In this study, we present the first reproducible, clinically relevant, murine model of African Trypanosomiasis optimized to characterize the functional changes and immunology of *T. brucei*-associated cardiac disease.

## Results

### *T. brucei* infection in mice results in infection dynamics and clinical progression which are comparable to HAT

Trypanosomiasis can be induced in mice by injecting viable *T. brucei* parasites into the tail vein of the mouse, resulting in a disease which is commonly used as a model for HAT (10, 31). To demonstrate that the mouse model recapitulates major features of natural infection, we infected adult male and female C57Bl/6J mice intravenously with 5-25 parasites of the *T. b. brucei* subspecies, a non-human infective subspecies of *T. brucei* which is most closely related to *T. b. rhodesiense* (4).

Following infection, parasitemia was measured every two days by hemocytometer. Parasitemia first became detectable in the bloodstream at 4-6 dpi, and the first peak of parasitemia occurred at 6-7 dpi (Fig 1A). At 8-10 dpi, the first trough of parasitemia occurred, when parasitemia reached undetectable levels. For the remainder of infection, parasitemia fluctuated and was highly variable between individual mice (Fig 1A).

**Figure 1.**
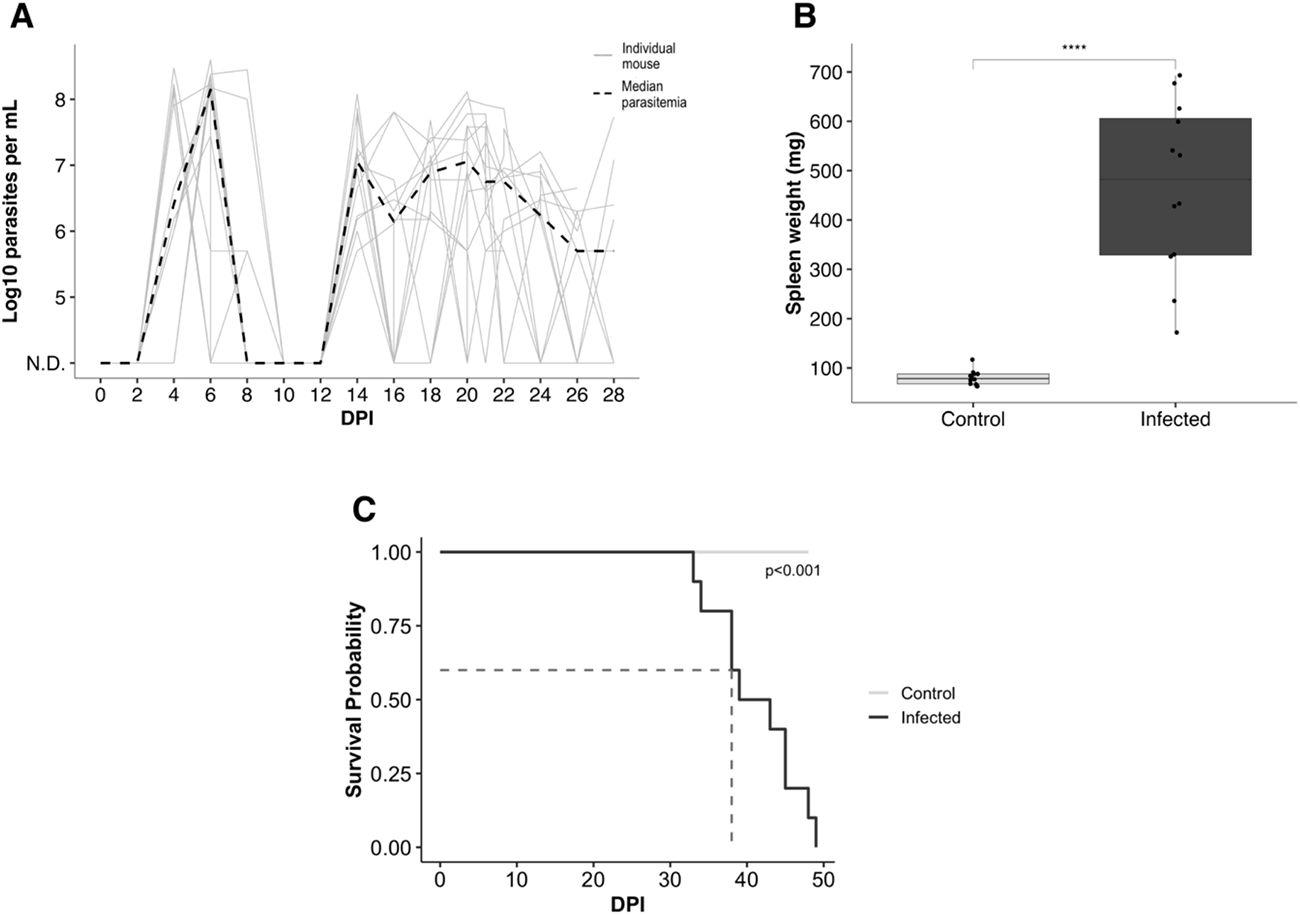
*T. brucei* infection causes waxing and waning waves of parasitemia and massive splenomegaly, with terminal endpoint reached within 50 dpi. A. Quantification of median and individual parasitemia in a representative experiment. Parasitemia first becomes detectable at 4-6 dpi. B. Weight of spleens of infected mice at 28 dpi(n=6) compared to uninfected, age-matched controls. Infected mice exhibit significant splenomegaly (Student’s t-test, p=3.7E-06). C. Kaplan-Meier survival curve of mice infected with T. brucei vs. uninfected control animals. Infected mice have a median survival time=39 dpi (p=4.32E-05).

*T. brucei* infection causes severe, multisystemic inflammation in natural hosts, with associated splenomegaly (2, 32). To confirm that *T. brucei* infection caused systemic inflammation in our model, mice were euthanized and the spleens were weighed at 28 dpi. Infected mice exhibited significant splenomegaly, with a mean splenic weight of 466 g, compared to 80.1 g in age-matched control mice (Fig 1B). Mice were euthanized when they reached 10^9^ parasites per mL of blood, lost >20% of body weight, or were assessed to be clinically terminal. Median survival time was 39 dpi, with the first mouse reaching terminal endpoint at 31 dpi and the last surviving until 49 dpi. Male mice exhibited a shorter survival time than female mice, although this difference was not statistically significant (Supp. Fig 2.1A).

The dynamics of *T. brucei* infection in our model were in general agreement with other published studies which have used similar mouse models and demonstrate that the mouse recapitulates natural infection (10, 31, 33).

### *T. brucei* infection induces elevation in NT-ProBNP and ECG abnormalities prior to a reduction in left ventricular ejection fraction

At 28 days postinfection (dpi)—a commonly used timepoint to represent chronic *T. brucei* infection—we evaluated cardiac biomarkers and function (31).

We measured cardiac NT-proBNP as a biomarker of cardiac dysfunction. Cardiac NT-proBNP is a sensitive marker of heart failure (34). Released from cardiomyocytes under conditions of excess stretch due to pressure or volume overload, NT-proBNP is elevated in HAT patients (16). Plasma NT-proBNP concentrations were significantly increased in infected mice at 28 dpi (Fig. 2A). NT-proBNP levels were slightly more elevated in male than in female mice, although this difference was not statistically significant (Supp. Fig. 2A).

**Figure 2.**
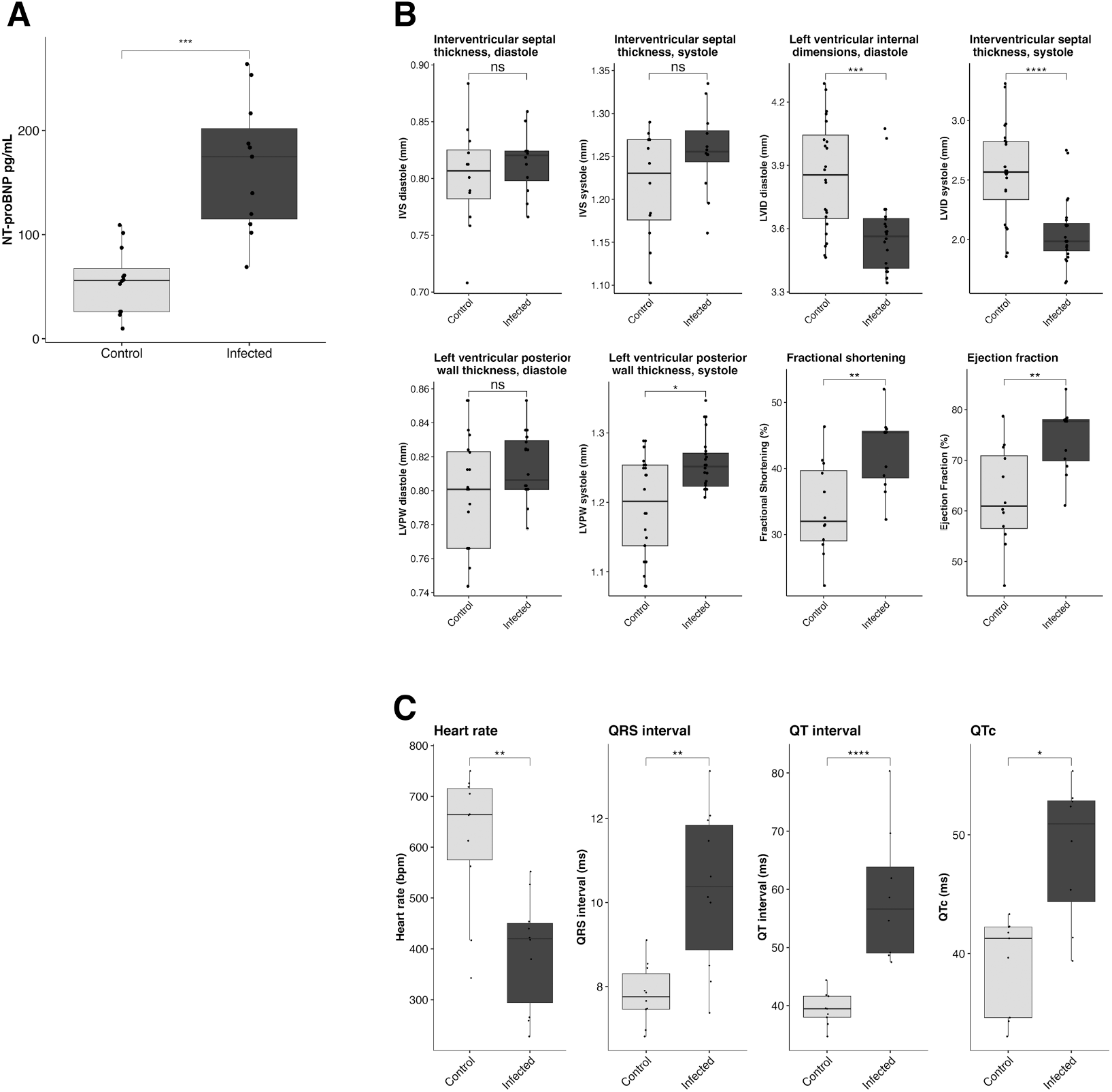
*T. brucei* infection causes elevated NT-proBNP, electrocardiographic abnormalities and heart failure with preserved ejection fraction. A. Measurement of plasma NT-proBNP at 28 dpi. Plasma NT-proBNP is significantly elevated at 28 dpi (n=12) compared to age-matched uninfected controls (Student’s t-test, p=0.00037). B. Echocardiographic measurements at 28 dpi. At 28 dpi, infected mice (n=12) exhibit significantly increased ejection fraction compared to uninfected age-matched controls, as measured by sedated echocardiography (Student’s t-test, p=0.0029). C. Electrocardiographic measurements at 28 dpi. At 28 dpi, infected mice (n=10) exhibit significantly decreased heart rate (HR, Student’s t-test, p= 0.00011), increased QRS interval, (Student’s t-test, p=0.00281), increased QT interval (Student’s t-test, p=0.00493), and increased QTc (Student’s t-test, p=0.01294) compared to uninfected age-matched controls.

We used echocardiography under sedation at 28 dpi to evaluate heart function. Infected mice had significantly increased ejection fraction and fractional shortening at 28 dpi (Fig 2B). Ejection fraction was slightly higher in infected males than in females, although this difference was not statistically significant (Supp. Fig. 2B). To evaluate the cardiac conduction system, we used ECG. Infected mice exhibited several ECG changes. As compared to uninfected controls, infected animals had a lower heart rate. A review of the EKG tracings showed that this was due both to a reduction in the intrinsic heart rate and to a significant burden of arrhythmias, including grouped beating and dropped beats, with significantly elevated QT intervals, including when QT interval was corrected for heart rate (QTc) (Fig. 2C). Overall, ECG findings were highly suggested of AV node dysfunction and intermittent high-degree atrioventricular block. ECG abnormalities were more exaggerated in infected males than females, although the differences were not statistically significant (Supp. Fig. 2C). All measured ECG parameters are shown in Table 1.

**Table 1:** Comparison of the mean values of all ECG parameters between infected animals and age-matched controls at 28 dpi (NS, P>0.05. *, P ≤0.05. **, P ≤0.01. ***,P ≤0.001) NS, P>0.05. *, P ≤0.05. **, P ≤0.01. ***,P ≤0.001

|  | Control | Infected | P value | Significance |
| --- | --- | --- | --- | --- |
| RR Interval (ms) | 104.98 | 166.77 | 0.00111 | ** |
| Heart Rate (BPM) | 616.18 | 394.40 | 0.00011 | *** |
| QRS Interval (ms) | 7.82 | 10.34 | 0.00281 | ** |
| QT Interval (ms) | 39.43 | 58.80 | 0.00493 | *** |
| QTc (ms) | 39.17 | 48.66 | 0.01294 | * |
| JT Interval (ms) | 31.77 | 48.25 | 0.00758 | ** |
| Tpeak Tend Interval (ms) | 28.97 | 42.24 | 0.00985 | ** |
| Q Amplitude (mV) | 0.01 | 0.02 | 0.42396 | NS |
| R Amplitude (mV) | 0.89 | 0.78 | 0.19872 | NS |
| S Amplitude (mV) | -0.52 | -0.30 | 0.00438 | ** |
| ST Height (mV) | 0.10 | 0.07 | 0.32718 | NS |
| T Amplitude (mV) | 0.19 | 0.11 | 0.00670 | ** |

NS, $P > 0.05$ . \*, $P \leq 0.05$ . \*\*, $P \leq 0.01$ . \*\*\*, $P \leq 0.001$

To further explore our observations, we repeated our functional assessments at 33 dpi, the latest timepoint at which mice can be experimentally manipulated before they begin to succumb to disease. Unsurprisingly, plasma NT-proBNP was again significantly elevated, with the mean NT-proBNP concentration (173.5 pg/mL) being similar to that observed at 28 dpi (165.4 pg/mL) (Fig. 3A). Due to the fragility of the mice at this timepoint, echocardiography was performed awake. Even so, 2 male mice and 1 female mouse had to be excluded due to reaching terminal endpoint before or during the echocardiographic evaluation. Interestingly, at 33 dpi mice exhibited a decreased ejection fraction, opposite to what was seen at 28 dpi (Fig. 3B). This could indicate cardiac decompensation as mice approach terminal status. It should be noted that, due to the fragility of mice at 33 dpi, echocardiography was performed awake, a technique that typically results in higher measured ejection fraction. Therefore, our findings indicate a significant drop in cardiac contractility between 28 dpi and 33 dpi.

**Figure 3.**
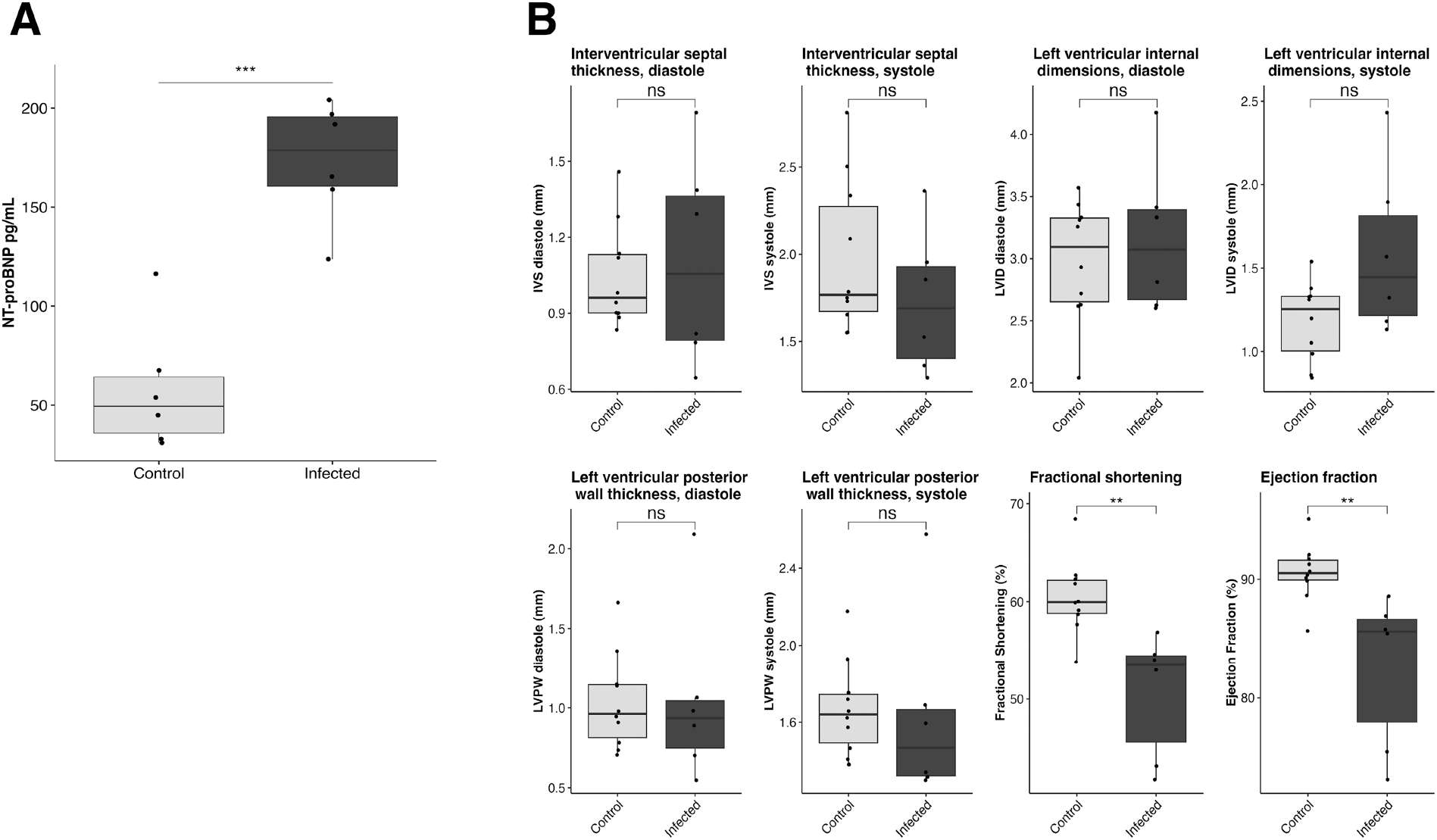
T. brucei infection causes elevated NT-proBNP and decreased heart function at terminal endpoint (33 dpi). A. Measurement of plasma NT-proBNP at 33 dpi. Plasma NT-proBNP is significantly elevated at 33 dpi in infected mice (n=6) compared to uninfected age-matched controls (Student’s t-test, p=2.8e-05). B. Echocardiographic measurements at 33 dpi. At 33 dpi, infected mice (n=6) exhibit significantly decreased ejection fraction compared to uninfected age-matched controls, as measured by awake echocardiography (Student’s t-test, p=0.011)

### *T. brucei* colonizes the myocardial interstitial space and causes myocarditis

Within the mammalian host, parasites live extracellularly and colonize multiple extravascular spaces, including the heart, ultimately causing fatal disease (13, 14). To confirm the presence of intracardiac parasites in our model and to demonstrate that these parasites are extravascular, we infected adult C57Bl/6J mice with genetically modified *T. brucei* parasites that constitutively express the tdTomato fluorescent protein (35)(35). This parasite line has previously been shown to exhibit similar infection dynamics as wild-type *T. brucei* parasites (13, 35). Mice were sacrificed at 14 dpi, a timepoint when extravascular parasite populations have been established and can be demonstrated on microscopy (10, 13). After sacrifice, we perfused mice to remove intravascular parasites, collected the heart, and performed immunofluorescence for CD31 to visualize the vasculature. *T. brucei* parasites were identified within the interstitial spaces of the heart. Intracardiac parasites localized separately from CD31-lined spaces, indicating that intracardiac parasites are extravascular (Fig 4, Supp. Fig 3).

**Figure 4.**
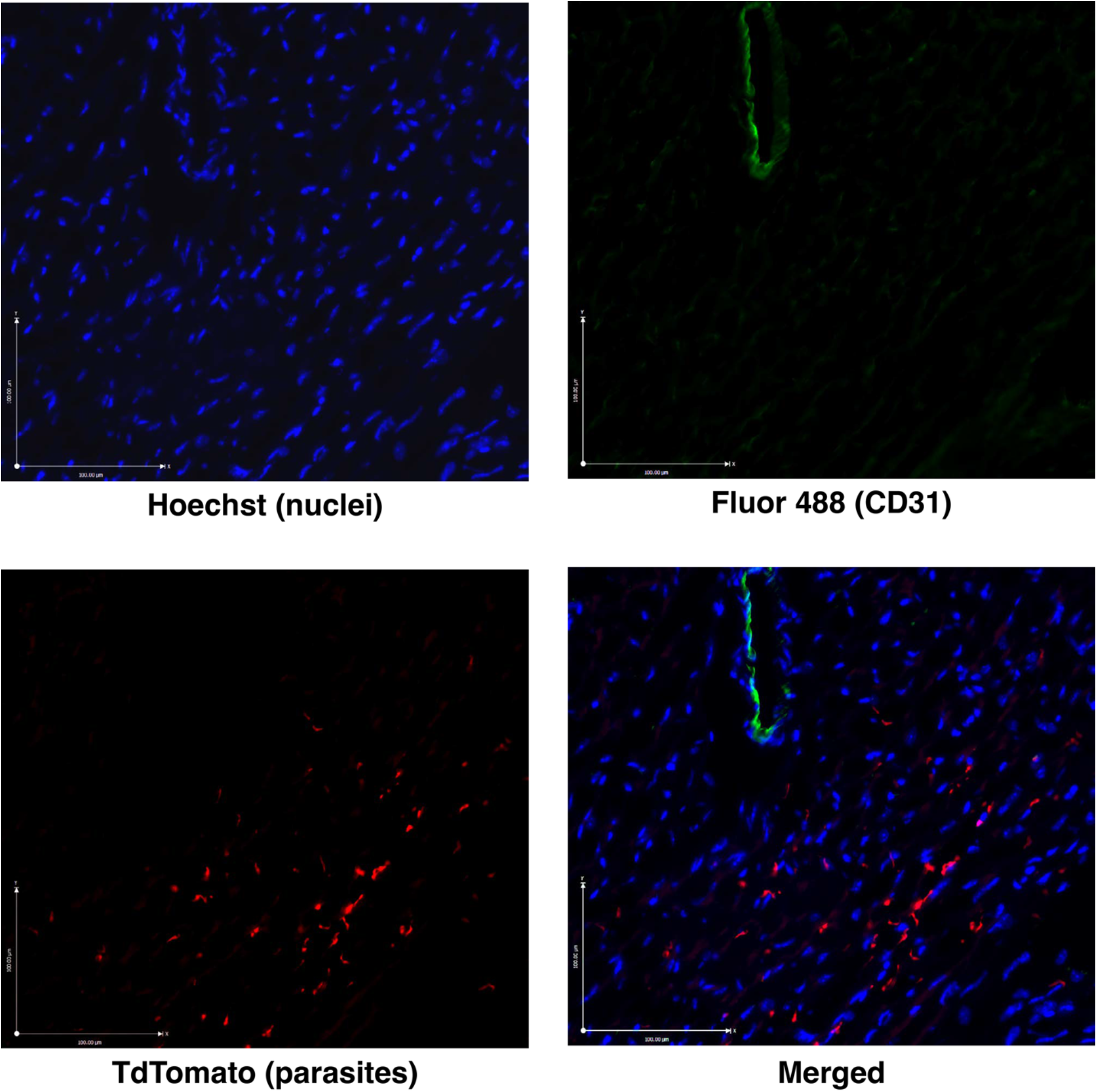
*T. brucei* parasites occupy extravascular spaces in the heart. Representative immunofluorescence microphotographs of the cardiac ventricle at 14 dpi at 20x magnification. *T. brucei* parasites (tdTomato—red) localize separately from CD31-lined vascular structures (Alexa Fluor 488—green), confirming the extravascular localization of parasites. Nuclei are stained with Hoechst (blue).

To evaluate cardiac pathology due to *T. brucei* infection, we performed histopathology on the hearts of 6 *T. brucei*-infected mice at 28 dpi. Of the 6 hearts evaluated, 5 met the Dallas Criteria for lymphocytic myocarditis, while the sixth exhibited borderline myocarditis (36). To further characterize histopathological changes, we used an established quantitative grading system to evaluate the severity of myocarditis (Table 2) (37). At 28 dpi, all mice exhibited myocarditis, with an average histological grade of 2, indicating moderate multifocal myocarditis (Fig 5A). Inflammation affected all layers of the heart, including endocardium, myocardium, and epicardium. On histopathology, observed cells included large numbers of lymphocytes, plasma cells, and macrophages. Necrosis was not a major feature. The severity of inflammation was most severe in the atria and at the atrioventricular (AV) junction (Fig 5B). The pattern and severity of inflammation observed were consistent with previous histopathology findings in humans and natural hosts (20, 22–25, 29, 38, 39). To evaluate cardiac fibrosis, longitudinal sections of the heart were stained with Masson’s Trichrome, and the percentage of collagen in the myocardium was quantified using the Automated Fibrosis Analysis Tool (AFAT) (40). At 28 dpi, infected mice had a higher percentage of collagen in the heart compared to uninfected mice, although this difference was not statistically significant (Fig 5C).

**Figure 5.**
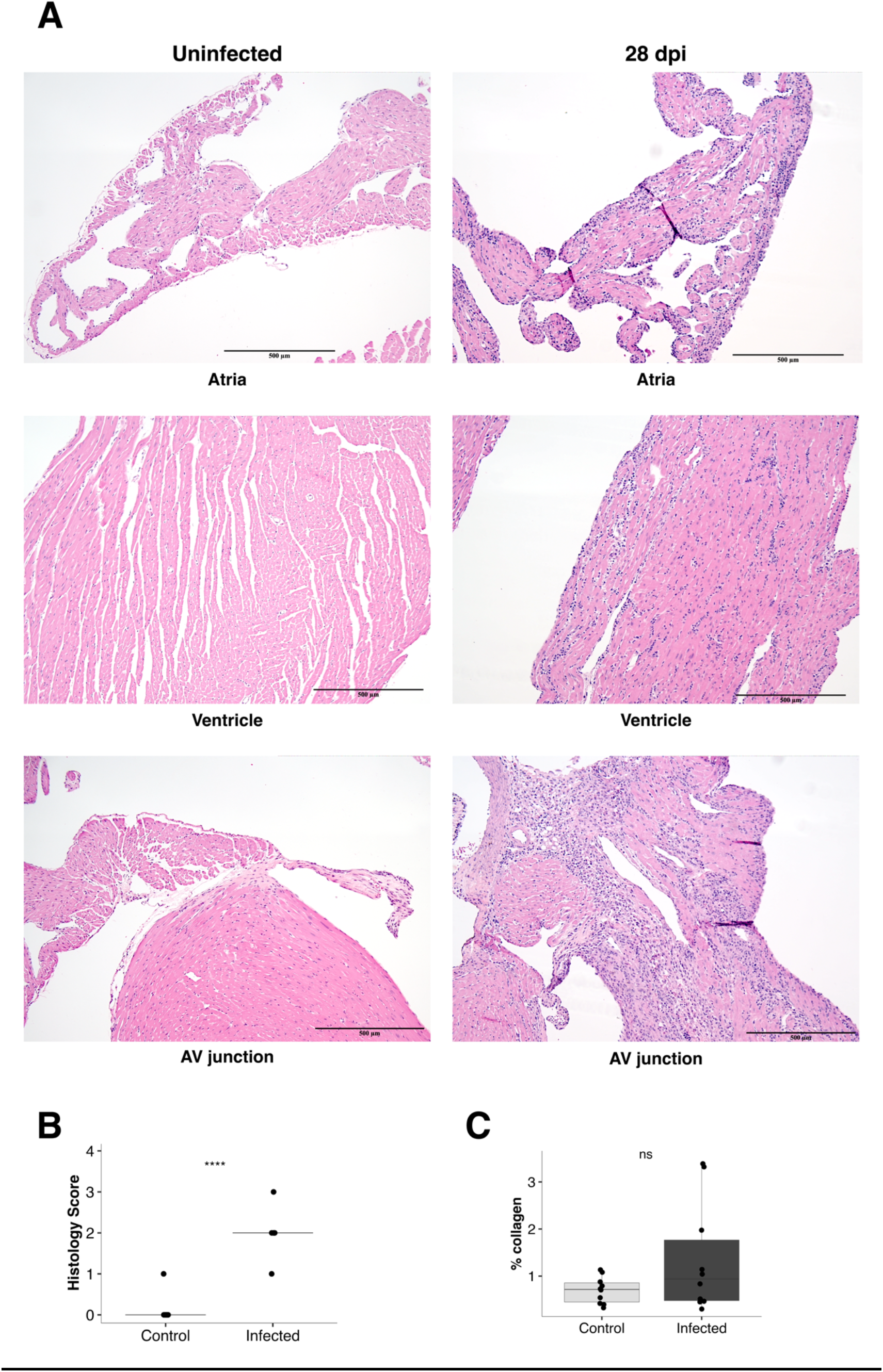
*T. brucei* infection causes myocarditis at 28 dpi. A. Representative brightfield microphotographs of the heart of infected mice at 28 dpi, compared to uninfected age-matched controls at 10x magnification. Inflammation is present in all layers and regions of the heart and is composed largely of mononuclear cells, primarily lymphocytes, plasma cells, and histiocytes. Inflammation is most severe at the atrioventricular junction. B. Histological grading of myocarditis (n=6) at 28 dpi, compared to equal numbers of age-matched controls. The average grade at 28 dpi is ‘2’, indicating multifocal moderate myocarditis without extensive associated necrosis. Histological grade is significantly higher in infected than uninfected mice (Student’s t test, p=1.5e-8). C. Measurement of the percentage of collagen in longitudinal sections of heart (n=6) at 28 dpi, compared to equal numbers of age-matched controls. There is no significant difference between infected and control mice (Student’s t test, p=0.31).

**Table 2:** Histological grading criteria.

| Grade | Description |
| --- | --- |
| 0 | No inflammatory infiltrates |
| 1 | Limited focal distribution of myocardial lesions |
| 2 | Multiple lesions with inflammatory infiltrates and some confluence |
| 3 | Multiple lesions with confluence and extensive necrosis |
| 4 | Lesions affecting most of the observed tissue |

### Immune cell populations in the heart are consistent with immune-mediated myocarditis

We had identified myocarditis as a consequence of *T. brucei* infection, and previous research has indicated that the host immune response may contribute to *T. brucei-*associated cardiac pathology (29). To better understand the intracardiac immune response to *T. brucei*, we performed flow cytometry at 28 dpi to characterize the immune cell population in the heart. Numbers of CD45+ cells/mg of tissue were markedly elevated in the hearts of all infected mice at 28 dpi, supporting our histopathological finding of myocarditis (Fig 6A).

**Figure 6.**
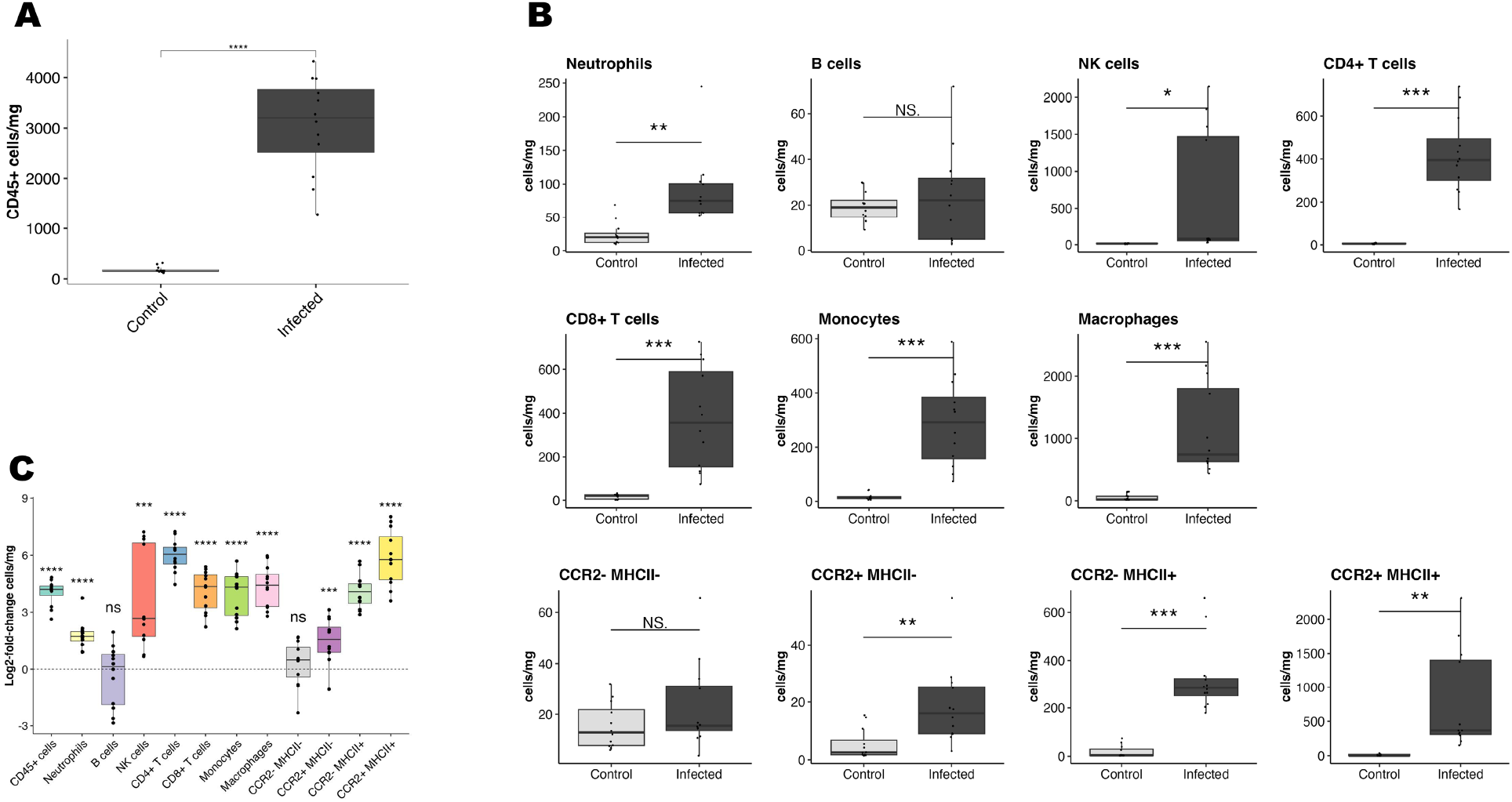
The intracardiac immune cell population during *T. brucei* infection is characteristic of immune-mediated myocarditis. A. Flow cytometric quantification of CD45+ cells per mg. At 28 dpi, CD45+ cells are markedly increased in the hearts of infected mice (n=12) compared to uninfected age-matched controls, supportive of myocarditis (Student’s t-test, p= 4.6e-7). B. Flow cytometric quantification of myeloid and lymphoid cell populations. Most immune cell populations are significantly increased in the hearts of infected mice at 28 dpi. Statistical comparisons were made using One-way Anova. *p<0.05, **p<0.01, ***p<0.001. C. Visualization of magnitude of changes in immune cell populations using the log2-fold change in cells/mg compared to uninfected age-matched control mice calculated based on the average cells/mg in the hearts of uninfected control mice.

Examining individual cell populations, most immune cell lineages were more abundant in *T. brucei* infected hearts than in age-matched controls (Fig 6B-C). In agreement with the functional data, most immune cell populations were slightly more elevated in males than in females, although the difference was not statistically significant for any cell lineage (Supp. Fig 2E-F).

Numbers of both CD4+ T cells and macrophages were significantly increased in infected hearts, which is unsurprising, since these cell types are necessary for an effective immune response to African trypanosomes (Fig 6B-C) (41). Of the different macrophage subsets, the CCR2^hi^MHC-II^hi^ population experienced the largest increase in numbers. This subset represents pro-inflammatory, monocyte-derived macrophages, indicating that *T. brucei* infection causes an influx of pro-inflammatory macrophages into the heart (42, 43). There was also a significant elevation of both CD8+ T cells and natural killer (NK) cells, even though cell-mediated immunity is relatively ineffective at responding to *T. brucei,* which is an extracellular parasite (42, 43).

## Discussion

*T. brucei*-associated cardiac disease is an underappreciated and understudied aspect of African trypanosomiasis. Here we report a reproducible murine model of *T. brucei* infection optimized to investigate acute and chronic *T. brucei*-associated cardiac disease. We demonstrate that this model recapitulates key aspects of *T. brucei*-associated cardiac disease in humans and natural hosts, including the presence of intracardiac parasites, myocarditis, EKG changes and elevated NT-proBNP. Using this model, we learned that *T. brucei* infection results in parasite colonization of the myocardial extracellular space and elevation of NT-proBNP (Figs 2-3). NT-proBNP elevation was initially present in the context of a preserved ejection fraction, but the infection eventually resulted in severe cardiac dysfunction as mice approached terminal endpoint (Figs 2-3). At a histological level, we found that myocardial colonization by the parasite triggered a severe myocarditis characterized by cardiomyocyte damage (Fig 5) and a multilineage inflammatory infiltrate (Fig 6). These findings advance our understanding of the host-pathogen interactions in the heart, suggesting that cardiac dysfunction associated with *T. brucei* infection is an immune-mediated event, potentially providing valuable information to inform the development of novel targeted therapies for animals and humans affected by African trypanosomiases.

Animal models have been used to study *T. brucei* infection since the early 1900s, and the first attempts to use a murine model to investigate *T. brucei*-associated cardiac disease were published in the late 1970s (15, 29, 30) . In these historical murine studies, investigators demonstrated cardiac lesions in infected mice and showed that immunosuppression was associated with a reduction in the size and frequency of inflammatory cardiac lesions (29).

However, despite their seminal nature, these studies were limited in the scope of their investigation, primarily focusing on histopathology and lacking more detailed functional or immunological assessments. Moreover, the studies were performed using infection protocols and parasite lines which were not standardized, thus making their methods challenging to replicate. Our present work expands significantly on those historical studies as it presents a reproducible method to generate cardiac damage and dysfunction in a murine model, demonstrably recapitulating multiple aspects of natural disease.

We have found that mice infected with *T. brucei* initially exhibit elevation in NT-proBNP and electrocardiographic abnormalities in the context of a preserved left ventricular ejection fraction. At 28 dpi, when we initially assessed BNP, the mice had clear histological evidence of myocarditis. NT-proBNP is elevated in response to an increase in intracardiac filling pressures (34). In the context of a normal ejection fraction and cardiac pathology, BNP elevation is highly suggestive of Heart Failure with Preserved Ejection fraction (HFpEF), a clinical condition characterized by reduced cardiac performance and symptoms of heart failure despite a seemingly normal cardiac contractility (44). However, BNP elevation in the context of a normal ejection fraction has also been described in patients with acute myocarditis, likely as a result of stiffening of cardiac walls triggered by the intracardiac immune response (45, 46). The potential overlap between myocarditis with preserved ejection fraction and elevated BNP and HFpEF is an area that has not been properly investigated so far in the clinical setting. However, taken together, our findings suggest that colonization of the cardiac interstitium by *T. brucei* might result in a stiffening of cardiac walls and elevation of cardiac pressures compatible with HFpEF, and that the myocarditic immune response triggered by the parasites likely plays a key role in this process. We hypothesize that as mice approach terminal endpoint, the cardiac pathology becomes severe enough to cause a decrease in ejection fraction. However, more definite conclusions require more detailed investigation into the functional impact of *T. brucei* infection at multiple timepoints.

At 28 dpi, electrocardiographic abnormalities included marked bradycardia, increased QT interval, grouped beating and dropped beats. These findings are in agreement with reports from HAT patients, in which increased QT interval is the most common findings (16, 17). AV blocks have also been reported in a subset of HAT patients, while autopsies have found inflammation of the sinoatrial node, suggesting a disruption of the cardiac conduction system (47). Interestingly, we did not observe premature ventricular contractions (PVCs), which have been observed in a rat model of African trypanosomiasis (26). However, the relatively short recording period in our study (1 minute per animal) may have hampered measurements of PVCs compared to the rat model, in which 30-minute ECG recordings were analyzed (26).

We used flow cytometry to investigate the intracardiac immune milieu during *T. brucei* infection. Previous studies in humans and large animal models have found that the major immune effectors during *T. brucei* infection are B cells, CD4+ T cells, and macrophages, which are all necessary for a protective anti-trypanosomal immune response (41). On the other hand, CD8+ T cells and natural killer (NK) cells are thought to be mediators of pathological inflammation in *T. brucei* infection, and their activation is associated with worse clinical outcomes (41, 48, 49). In agreement with our histopathological finding of myocarditis, we found that most intracardiac immune cell populations were elevated, especially CD4+ and CD8+ T cells, macrophages, and NK cells. The most highly elevated macrophage subset was the CCR2^hi^MHC-II^hi^ population, which represents pro-inflammatory, monocyte-derived macrophages (42, 43). Such pro-inflammatory macrophages are necessary for an effective immune response to *T. brucei* (41). However, increased activation of anti-inflammatory, alternatively-activated macrophages during chronic *T. brucei* infection is important for long-term survival in experimentally infected animals, due to their ability to attenuate pathological inflammation (47, 48). The large numbers of intracardiac CCR2^hi^MHC-II^hi^ in our model suggests that an imbalance between pro-inflammatory and anti-inflammatory macrophages could be a contributor to myocarditis in our model. The elevation of CD8+ T cells and NK cells in the heart are further suggestive of immunopathology as a contributor to cardiac damage. Increases in these cell populations are suggestive of a response to intracellular antigens, although *T. brucei* is entirely extracellular throughout its life cycle (48). Since CD8+ T cells are not a major feature of the immune response to extracellular pathogens, we hypothesize that their elevation in our model indicates a response to cardiac self-antigens, indicating an auto-immune component of the observed myocarditis. This hypothesis is corroborated by other evidence suggesting that *T. brucei*-associated organ damage has an auto-immune component. A recent study found that meningeal B cells produce high-affinity autoantibodies during *T. brucei* infection, suggesting that autoimmune responses may be a component of *T. brucei*-associated pathology in multiple tissues(50). Clinically, cardiomyopathy patients in Cameroon have been found to have high levels of both anti-trypanosomal and anti-heart antibodies, further indicating a connection between *T. brucei* infection and autoimmune heart disease (18).

Furthermore, we hypothesize that immunopathology in the heart during *T. brucei* infection follows a similar process to the pathogenesis of Chagas cardiomyopathy, caused by the New World trypanosome *Trypanosoma cruzi.* The leading theory to explain the pathogenesis of Chagas disease posits that parasite colonization of the heart induces inflammation and release of self-antigens, resulting in a breakdown of self-tolerance and immune destruction of cardiomyocytes (51). With chronicity, loss of cardiac muscle and replacement with scar tissue causes a collapse in cardiac function, ultimately ending in the characteristic Chagas cardiomyopathy (51). Our findings suggest that similar mechanisms might be at play during *T. brucei* infection.

There was no increase in the B cell population in the heart. This is somewhat surprising, since antibodies are the major mediators of immune clearance of *T. brucei*, and B cells are crucial for systemic anti-trypanosomal immunity (41). However, our findings are consistent with other tissue-specific flow cytometry experiments during *T. brucei* infection, which have identified no changes in number of B cells in the lungs or adipose bodies of *T. brucei*-infected mice (14, 33). In contrast, other models of myocarditis and heart failure show that intracardiac B cell populations are increased and play a major pathologic role during viral myocarditis and myocardial infarction (52–57). It is possible that B cell activation plays a role in *T. brucei*-associated cardiac disease, even if there is no expansion of local populations.

We found limited evidence of a sex-specific difference in *T. brucei*-associated cardiac disease. Male mice exhibited higher NT-proBNP at 33 dpi, higher numbers of intracardiac immune cells, and more severe echocardiographic and ECG changes than females, although these differences were not statistically significant. Our findings are in agreement with evidence that HAT is more severe in men than women, as well as recent findings which indicate that T. brucei causes more severe weight loss and adipose wasting in male rodents (58, 59). Our findings are also consistent with data indicating that myocarditis is more common in men than women (60). Further work will be needed to expand on these observations and dissect the mechanistic basis of sex differences in HAT.

Much about *T. brucei*-associated cardiac disease remains unknown, and there are serious gaps in our current knowledge, which this animal model could be used to investigate. Our findings raise concerns about long-term, post-treatment consequences of *T. brucei* infection. Most HAT patients are identified and treated, permanently clearing infection. However, the severity of the myocarditis reported by ourselves and others suggests that heart dysfunction might be only loosely connected with parasite presence. (20, 22–25, 29, 38, 39). If so, anti-parasitic treatment would not fully alleviate the cardiac symptoms of HAT, since self-directed inflammation could continue even with the removal of the inciting stimulus. Moreover, this suggests that immunomodulatory therapies could be considered for treatment of both humans and cattle to suppress inflammation that could contribute to persistent heart disease. There is a necessity to investigate the long-term and post-treatment consequences of *T. brucei*-associated myocarditis to determine whether HAT patients are at increased lifetime risk of heart failure. In addition, the effect of *T. brucei* subspecies on cardiac disease remains to be investigated. Our mouse model most closely recapitulates HAT due to *T. b. rhodesiense* infection. However, currently the majority of human cases are caused by *T. b. gambiense*, which often causes a disease of a more chronic character than *T. b. rhodesiense* (2). Although there is clinical evidence that all T. brucei subspecies cause cardiac disease, the differences in cardiac manifestations between subspecies are uncertain (16, 21, 61).

In summary, we describe the first reproducible, optimized, clinically relevant murine model of *T.* brucei-associated cardiac disease, a feature of African trypanosomiasis that is often overlooked in current textbooks and clinical resources. In the context of this model’s characterization, we produced evidence suggesting that *T. brucei*-associated cardiac damage is an immune-mediated event. Future studies with this model will be instrumental in identifying potential treatments for cardiac manifestations of African trypanosomiasis, a disease which continues to take an extensive human and economic toll in large regions of the world.

## Methods

### Sex as a biological variable

For all experiments, data from both male and female mice were collected and analyzed.

### Mouse and parasite strains and infection

Male and female C57Bl/6J mice (WT, strain# 000664 Jackson Laboratory) between 7-10 weeks old were each infected by intravenous tail vein injection with ∼5 pleiomorphic EATRO 1125 AnTat1.1E 90-13 *T. brucei brucei* parasites. To quantify parasitemia, blood was collected from the tail vein every two 2 days starting at 4 days post-infection (dpi), and parasites were counted via hemocytometer (Supplemental Figure 1). Blood (125 uL) was collected by a submandibular bleed at terminal endpoint (28 or 33 dpi). Unless otherwise described, mice were euthanized at terminal endpoint by intraperitoneal injection of ketamine/xylazine and perfused with 50 mL of PBS-Glucose (0.055M D-glucose) with heparin to remove intravascular parasites.

For immunofluorescence experiments, mice were infected intravenously with ∼5 AnTat1.1E chimeric triple reporter *T. brucei* parasites which express tdTomato (35). Parasitemia was measured as previously described. At 14 dpi, mice were sacrificed as previously described.

For flow cytometry experiments, mice were infected and parasitemia was measured as previously described. Mice were sacrificed at 28 dpi using CO2 euthanasia.

### Echocardiography

At 28 dpi, echocardiography was performed on sedated mice by a blinded investigator using a Vevo 2100 Ultrasound System (VisualSonics Inc.) as described previously (62).

At 33 dpi, echocardiography was performed on awake mice using a 40-MHz transducer (VisualSonics Inc.) as described previously (63).

### Electrocardiography

At 28 dpi, ECG was performed on unanesthetized mice as previously described (64). The entire procedure was performed in approximately 1 minute per mouse using a PowerLab data acquisition system (model ML866, ADInstruments, Colorado Springs, CO) and Animal Bio Amp (model ML136, ADInstruments). ECG data were analyzed using LabChart software (ADInstruments). The lead II tracings were used for analysis.

### Heart digestion

Heart digestion was performed using a technique previously (65). Briefly, after sacrifice, mice were perfused with 3 mL Hank’s Balanced Salt Solution (HBSS). 60 mg of heart tissue was placed in a 15 mL tube with 3 mL of HBSS. Digestion was performed for half an hour with 300 units of DNase I (Sigma-Aldrich), 625 units of collagenase II (Sigma-Aldrich), and 50 units of hyaluronidase II (Sigma-Aldrich). After digestion, the tubes were centrifuged at 250xg for 5 minutes at 4 °C, the supernatant was removed, and the pellet was resuspended in 5 mL ACK lysis buffer (Invitrogen) and incubated for 5 minutes at room temperature to remove red blood cells. Suspended cells were passed through a 50 μm filter and then prepared for flow cytometry.

### Flow cytometry

Flow cytometry was performed using a technique previously described (65). Cell suspensions were labeled with fluorescently conjugated antibodies (Supplemental Table 1). FACS was performed on an Aurora Spectral Flow Cytometer (Cytek) and analyses were performed using FlowJo version 10.8.1 (Becton Dickinson).

### NT-proBNP ELISA

At 28 and 33 dpi, plasma was collected from mice via cheek-bleeding. NT-proBNP plasma levels were measured using the LS-F34395 Mouse NT-proBNP Sandwich ELISA Kit (LSBio) according to the manufacturer’s instructions.

### Histopathology

For histopathology experiments, animals were infected and sacrificed at 28 dpi as previously described. Hearts were placed in 10% neutral buffered formalin and fixed at room temperature for 48 hours. The heart was cut in half longitudinally and sectioned by the Johns Hopkins Oncology Tissue and Imaging Services Core. To evaluate inflammation, hearts were stained with hematoxylin and eosin and imaged with 4x, 10x, and 40x objectives using a Nikon Eclipse E400 light microscope (Nikon). Cardiac lesions were graded by a blinded veterinary pathologist using a previously established both the Dallas Criteria and a semiquantitative scoring system for myocarditis in rodents (Table 2) (36, 37). Briefly, the severity and extent of lesions was graded using a five-point system from 0 to 4. A zero score indicates no infiltration. A ‘1’ score indicates very limited focal distribution of myocardial lesions. A ‘2’ score indicates multiple lesions. A ‘3’ score indicates multiple regions with confluence and extensive necrosis. A ‘4’ score indicates lesions affecting most of the observed tissue. Images were captured using an Excelis 4K camera (Unitron) and processed using ImageJ v1.53 image analysis software (NIH). All methods followed published guidelines for experimental pathology (66). To evaluate cardiac fibrosis, additional longitudinal sections of hearts were stained with Masson’s Trichrome and scanned at 200x magnification using an Axios Z.1 scanner (Zeiss). The percentage of collagen in each heart section was quantified using the AFAT (40).

### Immunofluorescence

For immunofluorescence experiments, 8-week-old female C57Bl/6J mice were infected intravenously with 5 triple marker *T. brucei* parasites (35). At 14 dpi, mice were euthanized and perfused as previously described. Heart was collected and fixed in 4% paraformaldehyde in PBS for 12 hours at 4 nC. Post-fixation, tissues were sectioned longitudinally, frozen embedded in O.C.T. Compound (Tissue-Tek), and cut using cryostat microtome into 10 μm sections by the Johns Hopkins Oncology Tissue and Imaging Services Core.

The following antibodies were applied to sections: rat anti-mouse CD-31 (SCBT) followed by goat anti-rat Alexa Fluor 488 (CST). Coverslips were mounted using ProLong Gold (Life technologies). Tissues were imaged with 4x, 10x and 20x objectives using a Nikon Eclipse 90i fluorescence microscope (Nikon) and X-Cite 120 fluorescent lamp (Excelitas) with an ORCA-ER digital CCD camera (Hammamatsu) and ImageJ v1.53 image analysis software (NIH). Image collection and analysis followed published guidelines for rigor and reproducibility in immunofluorescence (67).

### Statistics

Unless otherwise described, figures were generated and statistical analyses were performed using the Tidyverse package in R version 4.2.3 (68).

### Study approval

Animal experiments were performed with the approval of the IACUC of Johns Hopkins University under protocol MO22H163

## Supporting information

Supporting Data Values

## Data availability

Values for all data points in graphs are reported in the Supporting Data Values file.

## Author contributions

NPC, MRM and LA conceptualized the study. NPC, MDC, AKB, PS, and KG conducted experiments and acquired the data. NPC, MDC, AL, PS, AB, and KG analyzed the data. KG, LA, and MRM provided resources. NC wrote the original draft of the manuscript. NPC, AKB, LA, and MRM reviewed and edited the manuscript. MRM acquired funding. MRM and LA supervised the study.

## Acknowledgements

We would like to kindly thank Brice Rotureau for providing us with the chimeric triple reporter cell line.

The Aurora Flow Cytometer used in this study was funded by NIH Grant S10OD026859. We acknowledge the support of the JHU Ross Flow Cytometry Core.

Echocardiograms were performed by Nadan Wang of the JHU Small Animal Cardiovascular Phenotyping and Animal Model Core.

Histology support was provided by the JHU Oncology Tissue Services Core, funded by JHU Cancer Center Core Grant P30 CA006973.

This project was supported by the Johns Hopkins Specialized Center for Research Excellence in Sex Differences (U54AG062333) and The Foundation for Gender-Specific Medicine. NPC was supported by T32 OD011089.

