## Supplemental Materials for "A murine model of *Trypanosoma brucei-*induced myocarditis and cardiac dysfunction"

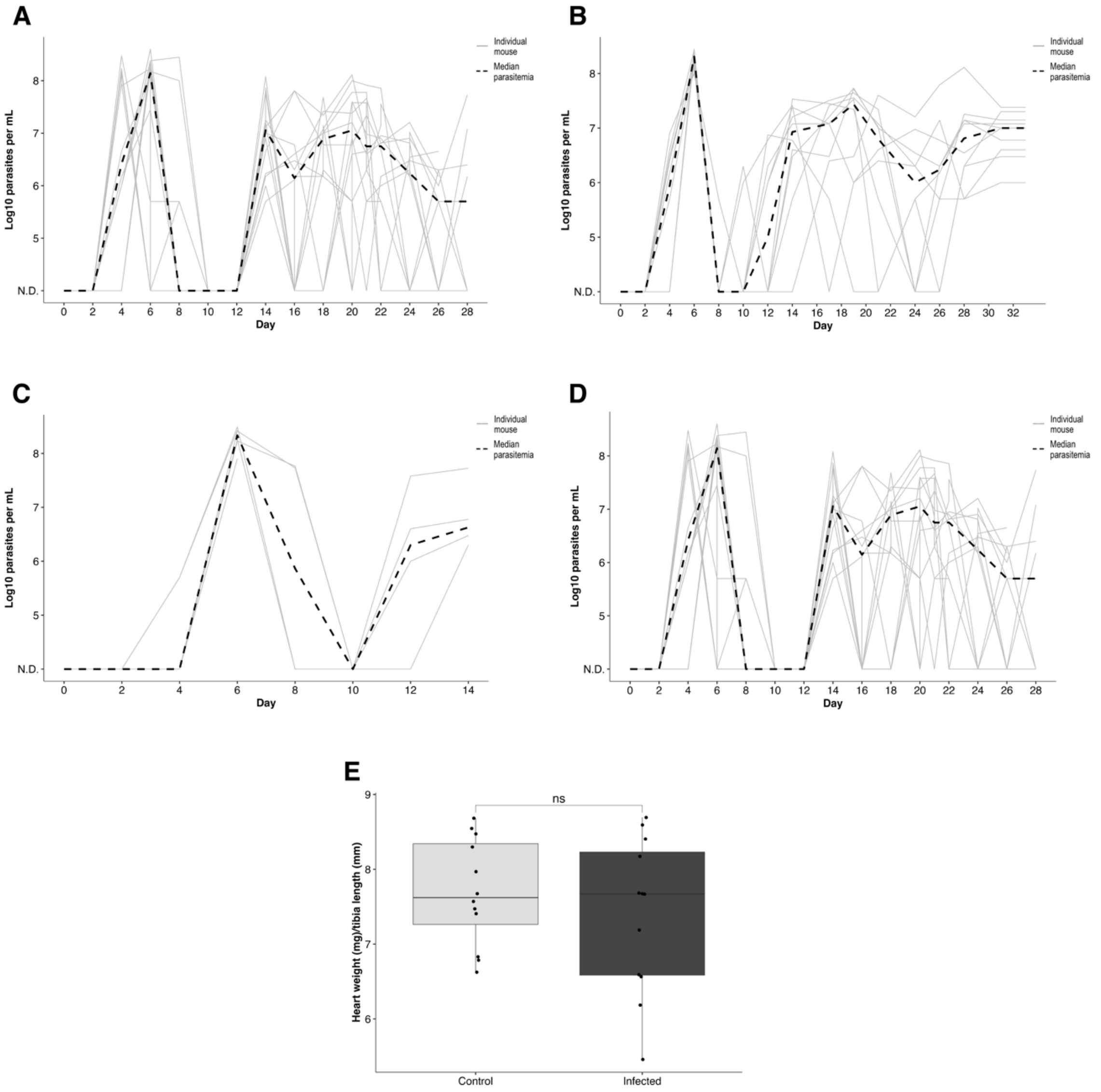


**Supplemental Figure 1:** Quantification of median and individual parasitemia for all experiments, heart weight at 28 dpi, and survival curve

A. Parasitemia of mice used for data collection at 28 dpi (NT-proBNP, echocardiography, histopathology)
B. Parasitemia of mice used for data collection at 33 dpi (NT-proBNP, echocardiography)
C. Parasitemia of mice used for data collection at 6 and 14 dpi (Immunofluorescence)
D. Parasitemia of mice used for survival curve and 28 dpi ECG
E. Heart weight in mg normalized to tibia length in mm of mice sacrificed at 28 dpi

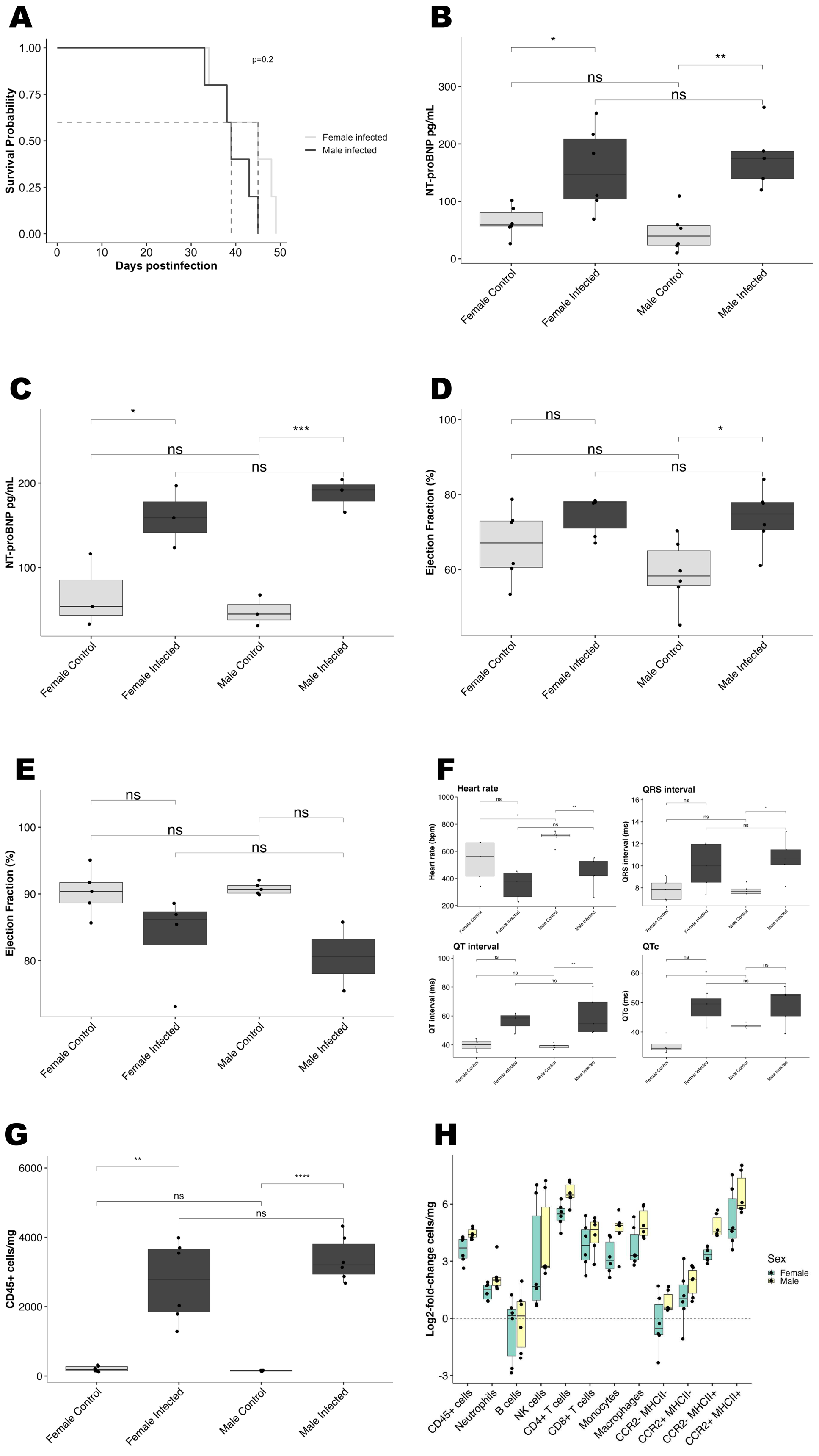
A
B
C
D
E
F
G
H
Survival Probability
Days postinfection
p=0.2
Female infected
Male infected
NT-proBNP pg/mL
Ejection Fraction (%)
CD45+ cells/mg
Female Control
Female Infected
Male Control
Male Infected
Heart rate
Heart rate (bpm)
QRS interval
QRS interval (ms)
QT interval
QT interval (ms)
QTc
QTc (ms)
Log2-fold-change cells/mg
Sex
Female
Male
CD45+ cells
Neutrophils
B cells
NK cells
CD4+ T cells
CD8+ T cells
Monocytes
Macrophages
CCR2- MHCII-
CCR2+ MHCII-
CCR2- MHCII+
CCR2+ MHCII+

**Supplemental Figure 2:** Experimental parameters of cardiac function and immunity, separated by sex

A. Survival curve separated by sex. Median survival time is 39 dpi for males and 45 dpi for females, p=0.2.
B. Plasma NT-proBNP measured at 28 dpi, compared between males and females (n=6 for each group)
C. Plasma NT-proBNP measured at 33 dpi (n=3 for each group)
D. Ejection fraction measured via sedated echocardiography at 28 dpi (n=6 for each group)
E. Ejection fraction measured via awake echocardiography at 33 dpi (n=4 for each group except Male Infected, for which n=2)
F. Electrocardiographic changes divided by sex (n=5 for each group)
G. Intracardiac CD45+ cells/mg measured via flow cytometry at 28 dpi (n=6 for each group
H. Log2-fold changes of all immune cell populations in infected mice at 28 dpi (n=6 for each group)

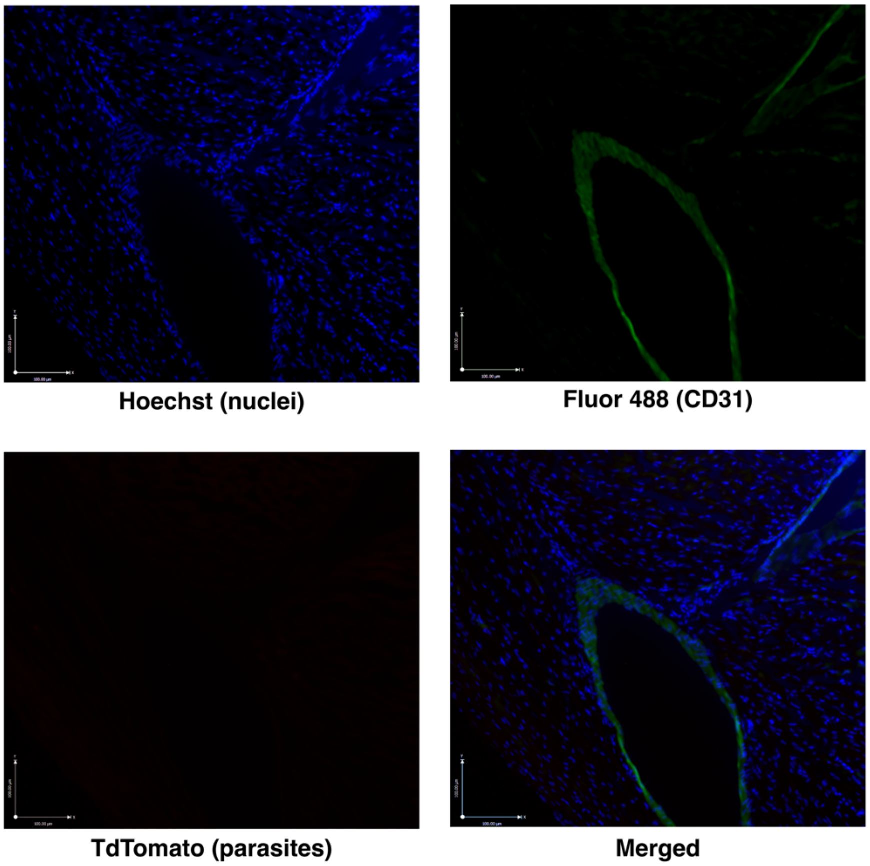

Hoechst (nuclei)

Fluor 488 (CD31)

TdTomato (parasites)

Merged

**Supplemental Figure 3:** Representative immunofluorescence microphotographs of the cardiac ventricle of an uninfected mouse at 200x magnification.

**Supplemental Figure 3**: Immunofluorescence of heart from uninfected mouse

**Supplemental Table 1:** Flow cytometry antibodies

| Antibody | Brand | Fluorophore |
|---|---|---|
| CD45 | Biolegend | PerCP/Cy5.5 |
| CD19 | Biolegend | BV421 |
| CD11b | BD horizon | AF700 |
| CD3 | Biolegend | APC Fire 810 |
| CD4 | Biolegend | SparkNIR |
| CD8 | Biolegend | BV785 |
| Ly6C | Biolegend | BV650 |
| Ly6G | Biolegend | APC Cy7 |
| CD64 | Biolegend | PECy7 |
| CCR2 | Biolegend | APC |
| MHCII | Biolegend | BV711 |
